# Decoding Cell Cycle Phase Variations in Cancer Hallmarks Across Breast Cancer Subgroups

**DOI:** 10.1101/2025.03.04.641383

**Authors:** Miguel Castresana-Aguirre, Alexios Matikas, Linda S. Lindström, Nicholas P. Tobin

## Abstract

Breast tumors have traditionally been categorized into four widely recognized subtypes. Recent single-cell advancements have, however, shown that individual tumors can contain more than one subtype, exhibiting their intrinsic heterogeneity. Relatedly, whilst the crucial role of cell cycle regulation in cancer development is widely acknowledged, the specific impact of cell cycle phases on oncogenic signaling and processes in each of these subtypes remains unresolved. Here, we examine the differences in transcriptomic cancer hallmarks across and within cell cycle phases and breast cancer subtypes in two large single-cell cohorts. We show that taking cell cycle phases into account identifies a broader pool of relevant cancer hallmark pathways while preserving the hallmarks found when cell cycle phases are ignored. Moreover, we identify FDA approved drugs that can target diverse hallmarks within specific phase-subtype cell combinations. This study highlights the critical role of cell cycle phases in advancing our understanding of breast cancer biology. The distinct biological mechanisms observed in each phase-subtype group suggest that targeting these hallmarks may provide a more precise and effective therapeutic strategy for breast cancer treatment.

## Introduction

Breast cancer accounts for approximately 31% of all female cancers and its incidence has increased by up to 0.5% annually since the early 2000s[1]. Traditional immunohistochemical (IHC) classification of breast cancer is based on the expression of the estrogen receptor (ER), progesterone receptor (PR) and human epidermal growth factor 2 (HER2) receptor. These biomarkers are used to divide breast tumors into luminal (ER+/PR+/−), HER2+ (HER2+, ER+/−, PR+/−) or triple negative (ER-, PR-, HER2-) subtypes. A further sub-division of the luminal subtype can be made with the proliferation marker Ki67 to classify tumors as Luminal A -like (ER+, PR+/−, Ki67 low) and Luminal B -like (ER+, PR+/−, Ki67 high). A genomic profiling approach can also be applied in order to derive breast cancer subtypes that are closely related but not identical to their immunohistochemical counterparts[2,3]. Applying the commercial version of Prediction Analysis of Microarray 50 (PAM50) signature to gene expression data from a breast tumor results in its classification into one of four main subtypes: Luminal A, Luminal B, Her2-enriched, Basal-like[4]. However, due to the intrinsic heterogeneity of breast tumors, classifying a whole tumor uniformly may not capture their biological complexity[5–8], a task that is better suited to single-cell RNA-sequencing (scRNA-seq) analyses[9]. scRNA-seq has enabled the analysis of the transcriptome at a single-cell level, providing a detailed view of the cellular landscape within tumor cell populations[10,11]. Recently, the bulk tumor PAM50 approach has been further developed in order to enable its application as a single-cell classifier (scPAM50) that can assign individual cancer epithelial cells to one of four breast cancer subtypes - Luminal A, Luminal B, Her2 and Basal[5]. In addition to this heterogeneity in single-cell molecular subtypes, a second seldom considered layer of complexity is that of tumor cell cycle phase.

Dysregulation of the cell cycle is a hallmark of cancer[12], uncontrolled cell proliferation, and tumor growth[13]. There are also clear differences in cell cycle activity levels between normal and tumor tissues[14], making the cell cycle a key target for cancer therapy[15–17]. Previous studies have shown that cells in different cell cycle phases show different transcriptomic profiles[18–21]. This is in line with the key concept that biological signaling pathways can be modulated by a cell’s progress through the cell cycle[22,23]. Relatedly, several studies have shown that understanding the interplay between cell cycle regulation and oncogenic signaling pathways is crucial for developing targeted therapies in breast cancer both at a bulk[24,25] and single-cell level[26–29]. To date however, studies examining the differences in pathways or cancer hallmarks as a cell proceeds through the cell cycle are lacking. To this end, we present a comprehensive single-cell analysis, in two large public cohorts, that aims to understand the critical role of the cell cycle in breast cancer. We determine expression differences in cancer hallmarks across the G0/G1, S and G2/M cell cycle phases and molecular subtypes before placing our results in the context of the potential implications for targeted therapy.

## Material and Methods

### Data processing and single-cell annotations

Two previously published single-cell breast cancer atlases were used for this study: a discovery cohort containing 100,064 cells from 26 patients[5] and a validation cohort containing 208,788 cells from 32 patients[30]. The whole genome scRNA-seq data with 20,224 annotated genes and basic clinico-pathological information was available for both cohorts. Cells were filtered out if fewer than 200 genes were detected in a cell or if their mitochondrial content was higher than 20%. The cohorts were log-normalized with a scaling factor of 10,000. Cell cycle phases (G0/G1, S, G2/M) were inferred using the Seurat package in R version 4.4.2 with predefined cell cycle markers[31]. In this study, we focused solely on cancer epithelial cells. This classification had already been made by the original authors in the discovery cohort and totaled 24,489 cancer epithelial cells. For the validation cohort, we inferred the cell types using the SingleR tool[32], having first trained and tested the model on the discovery cohort. This approach classified 95,401 cells as cancer epithelial in the validation cohort. scPAM50 subtypes were inferred for cancer epithelial cells using the scSubtype tool[5] to classify individual cells as Luminal A (LumA), Luminal B (LumB), HER2-enriched (Her2) and Basal.

### Differential Gene Expression (DGE) analysis and comparisons

As differential expression comparisons span cells across all tumors within discovery and validation cohorts (separately), addressing the confounding effect of tumor variation seems crucial. A recent comprehensive benchmark demonstrated however that for cohorts characterized by low depth (∼4%), high sparsity (∼90%), and significant batch effects, data integration could be detrimental and is not recommended[33]. Therefore, we selected the best performing approach from this benchmark which was a covariate model for breast cancer subtypes and cell cycle phases using LimmaTrend, limma R package v. 3.52.2[34], which is more suited to our cohort’s characteristics. Genes were then ranked for further analysis based on the moderated t-statistics.

In line with our aim to map the differences in cancer hallmarks across cell cycle phases taking breast cancer subtype into account, two distinct analysis approaches were adopted. In the first approach, we derived DGE lists by comparing cells from LumA, LumB, Her2 or Basal subtypes against each other within the same cell cycle phase (e.g., LumA G0/G1 cells vs. LumB G0/G1 + Her2 G0/G1 + Basal G0/G1 cells). In the second approach, we derived DGE lists by comparing cells from G0/G1, S or G2/M cell cycle phases against each other within the same scPAM50 subtype (e.g., LumA G0/G1 cells vs. LumA S + LumA G2/M). For clarity, we termed these analyses “*across subtype*” and “*within subtype*”, respectively and they are presented diagrammatically in Figure 1.

**Figure 1:**
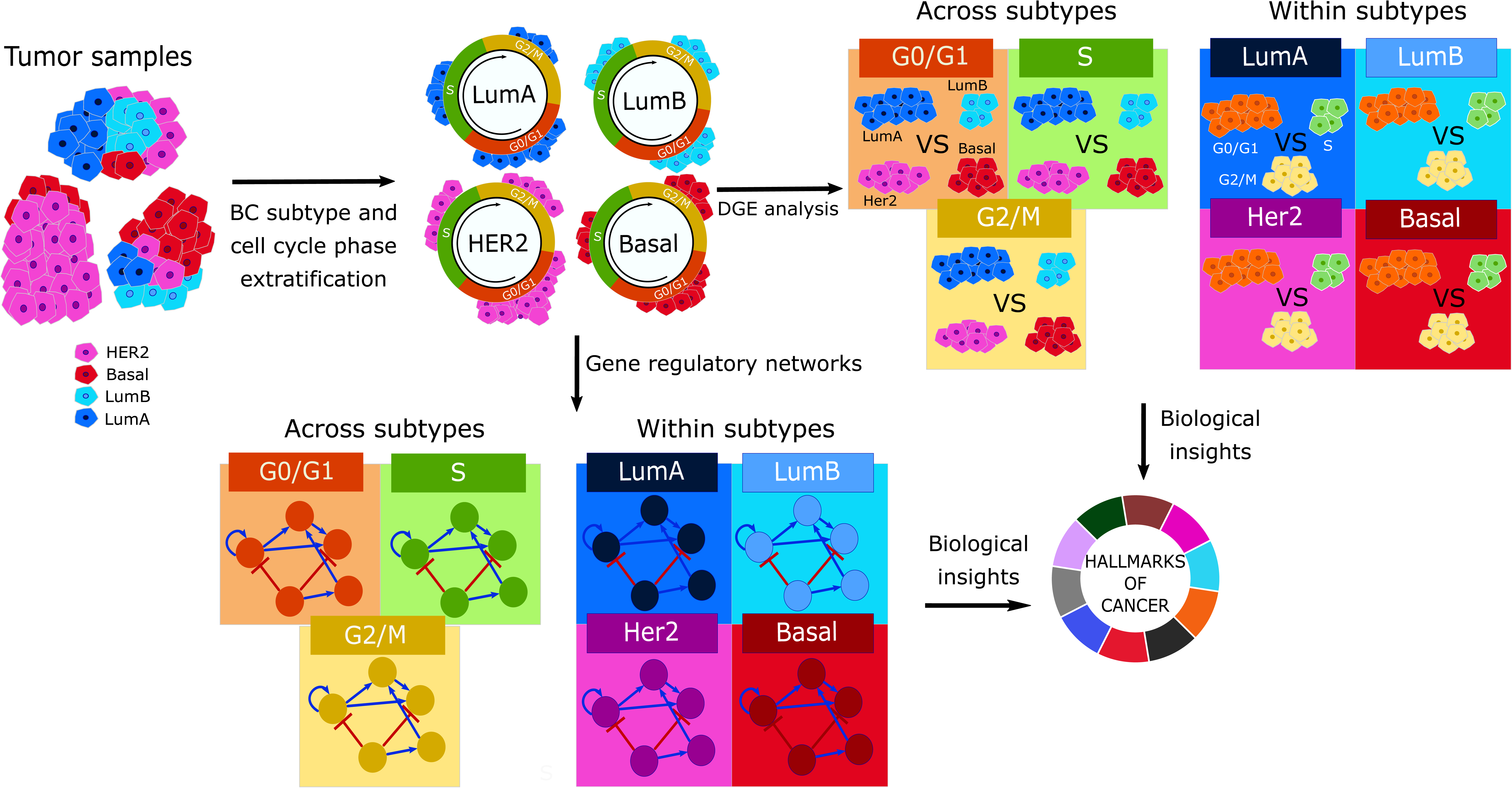
Overview of the study. Starting from patient tumor samples, we infer breast cancer subtype and cell cycle phase per cell. Then, in the *across subtype* comparison we perform differential expression analysis of breast cancer subtypes within cell cycle phases and infer a gene regulatory network per cell cycle phase. Finally, we look for biological insights by analyzing the hallmarks of cancer gene sets. In the *within subtype* comparison we perform differential expression analysis between cells of different cell cycle phases within breast cancer subtypes and we infer a gene regulatory network per subtype.

### Gene regulatory networks (GRNs)

GRNs were generated using pySCENIC, a python implementation of SCENIC[35], which resolves gene-to-gene interactions and gene co-expression modules. SCENIC identifies regulons that are composed of a transcription factor (TF) and its directly regulated genes by searching for over-represented DNA motifs near target genes. Thus, suggesting regulatory interactions and emphasizing biologically plausible relationships. The whole genome was used for GRN inference and the regulon activity in each cell was calculated using AUCell. For subgroup analyses, four GRNs were inferred - one for each breast cancer subtype, and for phase comparisons, three GRNs were delineated - one for each cell cycle phase. For example, the Luminal A GRN includes all cells of that subtype across all cell cycle phases. Regulons with regulon activity above the 95th percentile and with a Regulon Specificity Score (RSS) with a z-score > 1.5, indicating a tendency for expression in specific cell types (e.g., LumA G0/G1 cells), were selected for further analysis. The regulons that passed all filtering steps, showing sufficient relative activity and specificity, were used for pathway enrichment analysis.

### Pathway enrichment analysis

We focused on the hallmarks of cancer pathways from the Molecular Signatures Database (MSigDB) as the pivotal biological mechanisms underlying cancer development and progression. To gain insights into the relationship between the DGE lists and the MSigDB cancer hallmarks, we used the Fast Gene Set Enrichment Analysis (FGSEA) method[36], where pathways can be enriched (pathway genes tend to be overexpressed) or depleted (pathway genes tend to be underexpressed). For the regulons inferred from the GRNs, the overlap-based approach in the R package Clusterprofiler version 4.8.3[37] was utilized, analyzing if regulons significantly overlap with pathways of interest. To account for multiple testing, we applied the Benjamini-Hochberg procedure for correcting p-values[38]. Throughout the paper when we mention enriched or depleted pathways, they are statistically significant (FDR < 0.05).

### Drug targets and candidates

To identify potential drug targets and drug candidates of interest, we used two drug databases. The Comparative Toxicogenomics Database (CTD)[39], focusing on curated drug-targets relevant to *Homo sapiens* and specific to breast cancer, with an emphasis on mRNA-based drugs, and Drugbank[40] FDA approved drugs for *Homo sapiens*. The number of candidate drugs is 2106 and 1607 for CTD and Drugbank, respectively. Drug targets were selected according to the following criteria: TFs from the regulons identified in the GRNs that have associated drug candidates, pathways with statistical significance, regulons with an RSS with a z-score > 1.5, and regulons with activity above the 95th percentile.

### Statistical analysis

All analyses were performed using R version 4.4.2 and Python 3.9.19. To compare the proportion of cell cycle phases to scPAM50 subtypes we used a Chi-square test and a Cramer V test. To ensure robustness for pathway enrichment and drug targets-candidate analyses, results were only considered significant if the pathway/ drug was identified in the same phase and subtype in both cohorts.

## Results

### scPAM50 and cell cycle phase proportions

The scSubtype classifier was used to classify the single cancer epithelial cells of the discovery (N = 24,489) and validation (N = 95,401) cohorts into four breast cancer subtypes. In the discovery 6,809 (28%) individual cells were classified as LumA, 5,485 (22%) as LumB, 6,509 (27%) as Her2 and 5,686 (23%) as Basal (Supplementary Table 1, Discovery). In the validation cohort 13,247 (14%) cells were LumA, 36,378 (38%) LumB, 28,158 (30%) Her2, and 17,618 (18%) Basal (Supplementary Table 1, Validation). We first compared the proportions of scPAM50 subtype calls within individual tumors to their corresponding IHC classifications. Generally, the highest proportion of single-cell subtype calls within each tumor aligns with its IHC across both discovery and validation cohorts (Figure 2a and b, respectively). For example, the first tumor CID3941 (Figure 2a) is classified as ER+ by IHC, and the most frequent/highest proportion of single cells are classified as LumA (53%) by the scSubtype predictor. Luminal A tumours have been previously shown to be predominantly ER+ [3]. In the validation cohort, LumB was more frequently identified as the highest proportion of single-cell subtypes within individual tumours and we observe an overall matching between IHC and most frequent sc-subtypes for ER+, HER2+ and TNBC tumors.

**Figure 2:**
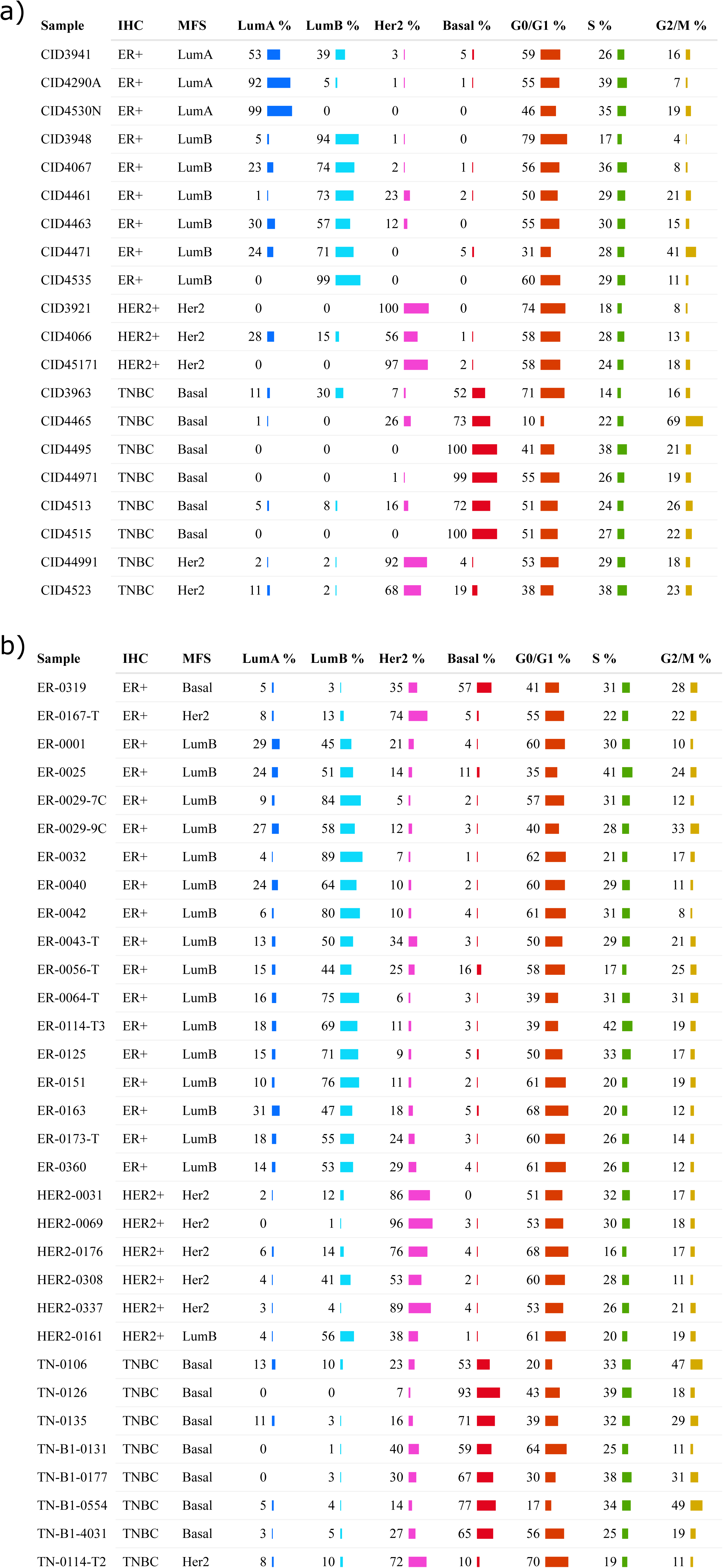
a) Discovery cohort with 20 samples and 24,489 cells. b) Validation cohort with 32 samples and 96,128 cells. MFS: Most Frequent Subtype.

In addition to subtypes, single cells were also categorized into three main cell cycle phases G0/G1, S, and G2/M. Overall in the discovery cohort, 12,992 (53%) were found to be in the G0/G1 phase, 7,660 (31%) in S, and 3,837 (16%) in G2/M (Supplementary Table 1, Discovery). In the validation cohort 50,136 (53%) of cells were in G0/G1, 26,647 (28%) in S, and 18,618 (20%) in G2/M (Supplementary Table 1, Validation). We also assessed single cell - cell cycle phases in relation to scPAM50 subtypes and found a statistically significant association in both discovery and validation cohorts (Chi-square test, P < 0.0001, Supplementary Table 1). However, Cramér’s V values of 0.10 and 0.06, respectively, suggest that the association is weak. This implies that while the relationship is statistically significant, it only accounts for a small proportion of the variability between the cell cycle phases and scPAM50 subtypes.

### Differential gene expression analysis identifies altered subtype and cell cycle phase specific cancer hallmark pathways

#### Gene set enrichment: Across subtype cancer hallmark pathway analysis

Next, we aimed to determine if taking cell cycle phases into account resulted in the identification of additional differentially expressed cancer hallmarks relative to not taking phases into account. For this, we conducted bulk comparisons *across subtypes* without accounting for cell cycle phase (e.g. all single LumA cells from all tumors versus all single LumB, Her2, and Basal cells). The same analysis was then repeated, comparing each subtype to the remaining three, but this time the same cells were separated into both single-cell subtypes and G0/G1, S and G2/M cell cycle phases (e.g. LumA G0/G1 vs. LumB G0/G1 + Her2 G0/G1 + Basal G0/G1). In total, bulk analysis identified 55 significantly differentially expressed cancer hallmark pathways when stratified by subtypes alone and phase-specific stratification identified 71. As depicted in the Venn diagram in Figure 3a, all pathways identified in bulk analysis are also present in phase-specific instances. This indicates that unique biological information emerges from subtype comparisons that incorporate cell cycle phase distinctions.

**Figure 3:**
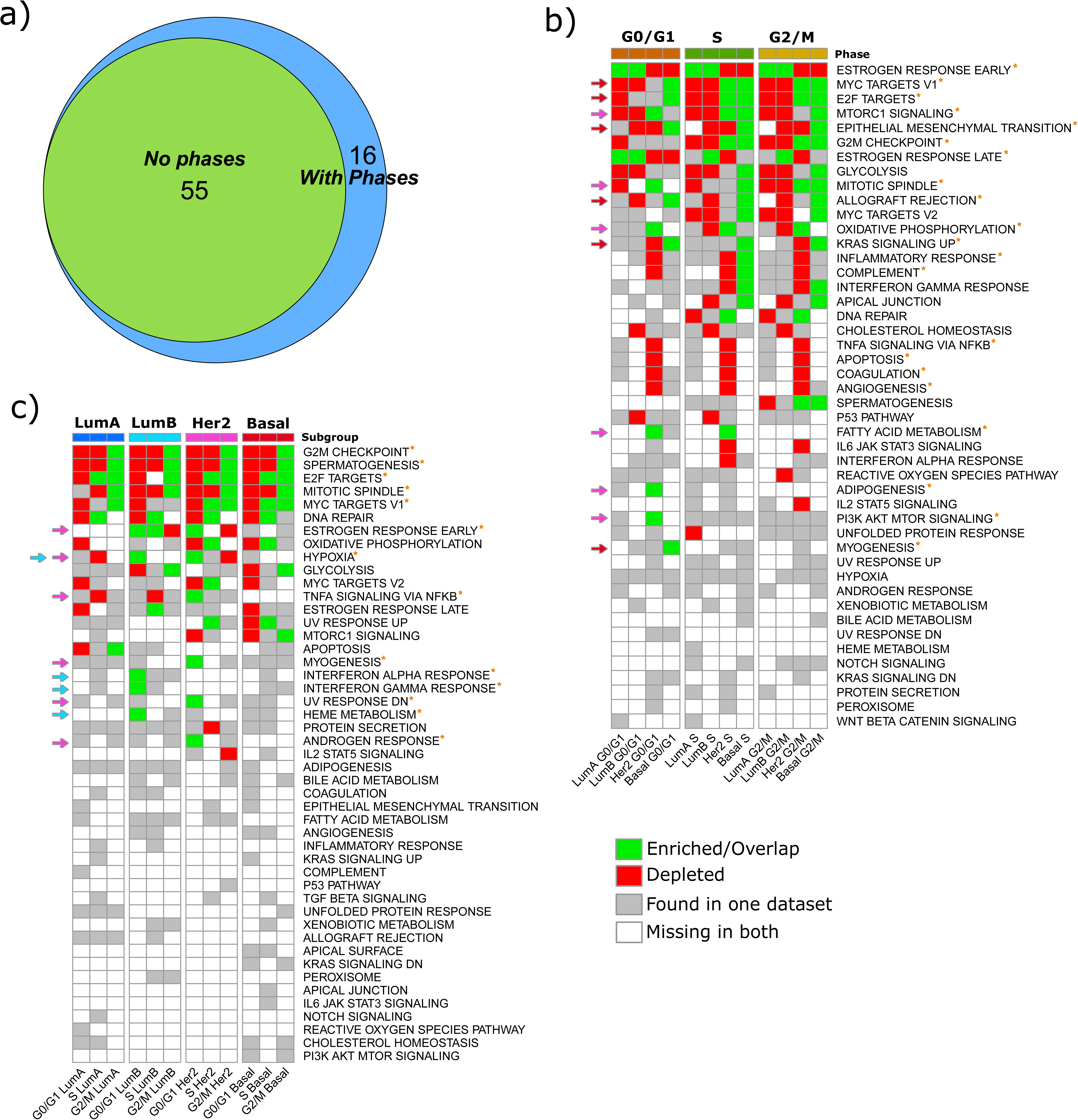
Differential gene expression analysis for hallmarks of cancer using FGSEA. Red means that the pathway is depleted in both discovery and validation cohorts, green means that it is enriched in both and gray means that there is no consensus between both cohorts. a) Overlap between the statistically significant pathways with and without taking cell cycle phases into account. b) *Across subtype* comparison, cells in the same cell cycle phase but in different breast cancer subtypes are compared (e.g. Luminal A against Luminal B, Her2, and Basal in G0/G1 phase). c) *Within subtypes* comparison, cells in the same breast cancer subtype are compared (e.g. Luminal A in G0/G1 against Luminal A in S and Luminal A in G2/M).

Focusing specifically on these 71 pathway-subtype-phase findings we observed biological signaling commonalities regardless of the individual cell cycle phase being analyzed. Specifically, Her2 and Basal cells had more enriched pathways relative to LumA/B cells in all phases (Figure 3b, compare subtype columns in G0/G1, S and G2/M phases, green = enriched, red = depleted). The ESTROGEN_RESPONSE_EARLY pathway was enriched in LumA/B and depleted in Her2 and Basal subtype cells in all phases. MYC_TARGETS_V1, E2F_TARGETS and G2M_CHECKPOINT hallmarks were depleted in LumA/B and enriched in Her2 and Basal subtype cells in S and G2M phases. Her2 cells exhibit a significant enrichment of OXIDATIVE_PHOSPHORYLATION, and depletion of INFLAMMATORY_RESPONSE, COMPLETEMENT, TNFA_SIGNALING_VIA_NFKB, APOPTOSIS, COAGULATION, and ANGIOGENESIS pathways in all cell cycle phases whilst Basal cells show the highest enrichment of pathways related to EPITHELIAL_MESENCHYMAL_TRANSITION, ALLOGRAFT_REJECTION, and KRAS_SIGNALING_UP in all phases (hallmarks mentioned above are denoted with a “*” in Figure 3b).

As G0/G1 cells likely contain a subpopulation of cells that may be treatment resistant[41,42], it is pertinent to highlight enriched hallmarks that could form the basis of a targeted treatment in this phase. Aside from the enrichment of estrogen response pathways noted above for LumA and B cells, we also found that Her2 cells show enrichment for MTORC1_SIGNALING, MITOTIC_SPINDLE, OXIDATIVE_PHOSPHORYLATION, FATTY_ACID_METABOLISM, ADIPOGENESIS and PI3K_AKT_MTOR_SIGNALING hallmarks (Figure 3b, see pink arrows on the left-hand side). Basal cells showed enrichment of MYC_TARGETS_V1, E2F_TARGETS, EPITHELIAL_MESENCHYMAL_TRANSITION, ALLOGRAFT_REJECTION, KRAS_SIGNALING_UP and MYOGENESIS (Figure 3b, red arrows).

#### Gene set enrichment: Within subtype cancer hallmark pathway analysis

In our initial analyses we studied DGE focused on the differences *across subtypes* when considering the same cell cycle phase. Next, we studied DGE by comparing each cell cycle phase to the remaining two *within* the same subtype.

This analysis highlighted biological signaling commonalities regardless of the individual subtypes being analyzed. G2M_CHECKPOINT, SPERMATOGENESIS and MITOTIC_SPINDLE were all depleted in G0/G1 and S-phases and enriched in G2/M cells for all four subtypes except for MITOTIC_SPINDLE for LumA G0/G1. Similarly, the MYC_TARGETS_V1 hallmark was depleted in G0/G1 and enriched in G2/M cells for LumA, Her2, and Basal subtypes but interestingly it was not enriched for LumB G2/M (Figure 3c). In addition, we found that whilst a pathway can show enrichment *within* the same subtype when comparing its cell cycle phases to each other it can still be depleted when comparing *across* subtypes. The E2F_TARGETS pathway provides a good example of this. When examining changes between cell cycle phases *within* LumA cells, we found this pathway to be depleted in the G0/G1 phase relative to the S and G2/M phases but enriched in the S (vs. G0/G1 + G2/M) and G2/M (vs. G0/G1 and S) phases (Figure 3c, see “E2F_TARGETS” in the heatmap). When comparing *across* subtypes however we found the pathway to be depleted in LumA cells in all cell cycle phases relative to LumB, Her2 and Basal cells (Figure 3b, see “E2F_TARGETS”). Put simply, while a pathway can be depleted in one subtype when compared *across* all other subtypes, it can still show differences in depletion and enrichment when comparing cell cycle phases *within* the same subtype.

Finally, we again highlight hallmarks that are enriched in the G0/G1 phase. We found that HYPOXIA is enriched in this phase in LumB and Her2 cells. In LumB cells alone we found INTERFERON_ALPHA, INTERFERON_GAMMA, and HEME_METABOLISM enriched. Finally, in Her2 cells alone we found ESTROGEN_RESPONSE_EARLY, TNFA_SIGNALING_VIA_NFKB, MYOGENESIS, UV_RESPONSE_DN, and ANDROGEN_RESPONSE enriched. (Figure 3c, see colored arrows on the left-hand side in combination with the subtype group at the top of the plot. Light blue arrows = LumB cells, pink = Her2 cells).

### GRN analysis

Differential gene expression analysis identifies genes with statistically significant expression variations across different conditions or groups. GRNs is a complementary methodology that aims to disentangle the complex interplay of these genes within networks. Thus, while DGE pinpoints genes of interest, GRNs provide the contextual and mechanistic insights necessary for developing targeted therapies.

#### GRN: Across subtype cancer hallmark pathway analysis

Similarly to our FGSEA analysis, we initially conducted bulk comparisons *across subtypes* without accounting for cell cycle phase. In total, bulk analysis identified 27 significantly differentially expressed cancer hallmark pathways when stratified by subtype alone, while phase-specific stratification identified 72. Of these, 24 subtype-pathway hits overlapped with bulk analysis, whereas 3 and 48 subtype-pathway combinations were unique to bulk and phase-specific analyses, respectively (Figure 4a). This indicates again, as DGE analysis did, that unique biological information emerges from subtype comparisons that incorporate cell cycle phase distinctions.

**Figure 4:**
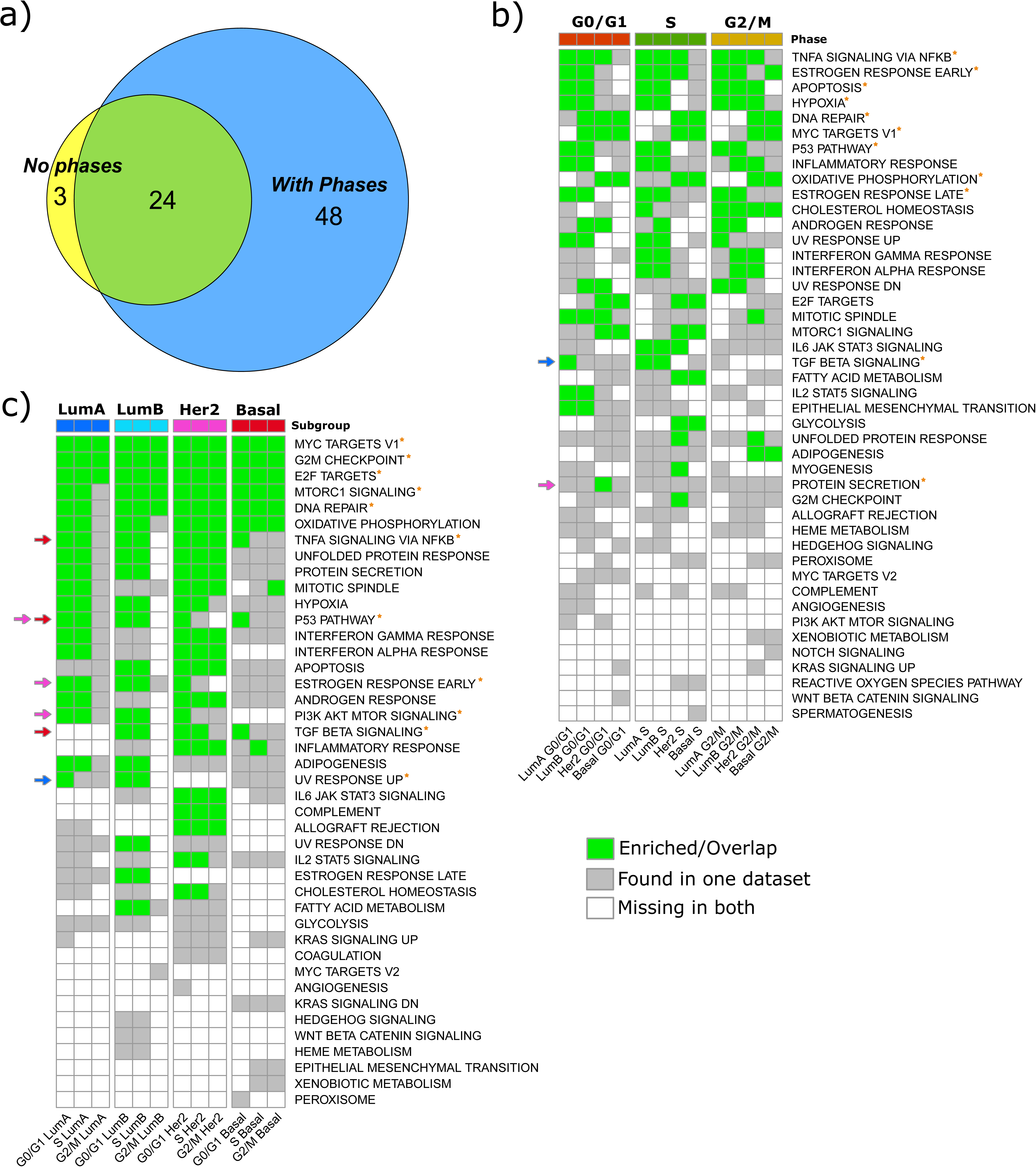
Gene regulatory networks and corresponding regulon enrichment for hallmarks of cancer using clusterProfiler. Green means that the pathway is enriched in both discovery and validation. a) Overlap between the statistically significant pathways with and without taking cell cycle phases into account. b) *Across subtype* comparison, a GRN is inferred for cells in the same cell cycle phase but in different breast cancer subtypes (e.g. all cells in G0/G1 phase for all breast cancer subtypes). c) *Within subtype* comparison, a GRN is inferred for cells in the same breast cancer subtype but in different cell cycle phases (e.g. all cells in Luminal A for all cell cycle phases).

Next, we again examined the biological signaling commonalities found across all cell cycle phases (Fig 4b). Note that in our GRN analysis, we used Geneset Enrichment Analysis (GEA) to determine pathways that statistically overlap with the regulons of interest. TNFA_SIGNALING_VIA_NFKB, ESTROGEN_RESPONSE_EARLY, APOPTOSIS, HYPOXIA, P53_PATHWAY, and ESTROGEN_RESPONSE_LATE hallmarks were statistically significant in both LumA and LumB cells in all cell cycle phases. For Her2 and Basal cells DNA_REPAIR, MYC_TARGETS_V1 and OXIDATIVE_PHOSPHORYLATION hallmarks were statistically significant in all phases.

Focusing again on the G0/G1 phase, examples of pathways unique to a subtype in G0/G1 included TGF_BETA_SIGNALING in LumA cells and PROTEIN_SECRETION in Her2 cells.

#### GRN: Within subtype cancer hallmark pathway analysis

The *within* subtype GRN analysis showed for example that the MYC_TARGETS_V1, G2M_CHECKPOINT, and E2F_TARGETS pathways were significant in all subtypes for all cell cycle phases. MTORC1_SIGNALING and DNA_REPAIR were significant in all phases in LumB, Her2 and Basals cells but not in LumA G2/M (Figure 4c).

As previously, we again highlight hallmarks that are enriched in the G0/G1 phase. We found P53_PATHWAY to be statistically significant for Her2 and Basal cells in this phase. In LumA cells we found UV_RESPONSE_UP. In Her2 cells alone we found ESTROGEN_RESPONSE_EARLY and PI3K_AKT_MTOR_SIGNALING. Finally, in Basal cells alone we found TNFA_SIGNALING_VIA_NFKB and TGF_BETA_SIGNALING (Figure 4c, see colored arrows on the left-hand side in combination with subtype groups at the top of the plot).

### Drug targets and drug candidates

Finally, we sought to identify potential drugs from both *across* and *within* GRN analyses for specific subtype-phase enriched pathways. Our analysis found 5 drugs and 7 chemicals from the Drugbank and CTD databases, respectively that target a transcription factor-subtype-phase-pathway combination. As multiple drugs/chemicals can target the same combination we provide examples here of FDA approved drugs only for each hallmark. The complete list of all drugs and chemicals is presented in Supplementary Table 2 and all combinations are shown in the form of multipartite network plots in Figure 5 (for Drugbank) and Supplementary Figure 1 (for CTD). The Drugbank drugs we identified targeted two main TFs (FOS and JUN) in LumA, LumB and Her2 cells, whereas in Basal cells only JUN was identified. Specifically, in the G0/G1 phase, drugs were found for the TNFA_SIGNALING_VIA_NFKB hallmark (e.g. Adapalene or Vinblastine) in all four breast cancer subtypes, for the ESTROGEN_RESPONSE_EARLY and ESTROGEN_RESPONSE_LATE hallmarks (e.g. Nandrolone decanoate) in LumA and LumB cells, for the P53_PATHWAY (e.g., Irbesartan) in LumB and Her2 cells and for UV_RESPONSE_UP (e.g., Adapalene) in LumB cells only. In the S phase, 4 targetable pathways were found for both LumA and B cells: APOPTOSIS, P53_PATHWAY, TNFA_SIGNALING_VIA_NFKB, and HYPOXIA (e.g. all targetable with the drug Irbesartan). Additionally, UV_RESPONSE_UP (e.g. Vinblastine) can be targeted in S phase LumB cells as can TNFA_SIGNALING_VIA_NFKB pathway (e.g. Irbesartan) in Her2 cells. Finally, for the G2/M phase, TNFA_SIGNALING_VIA_NFKB and APOPTOSIS (e.g. Nandrolone decanoate) were found to have significant drugs for both LumA and LumB cells. The CTD chemicals we identified targeted three main TFs (FOS, JUN and FOXM1) in all subtype cells and all chemicals along with the specific transcription factor-subtype-phase-pathway combinations they target are shown in Supplementary Table 2 and Supplementary Figure 1. Taken together, these results demonstrate the possibility of identifying and targeting key hallmarks within specific subtype-phase combinations. This may enhance future precision oncology initiatives and promote the use of combinational or sequential therapies for patient treatment (an illustrative example is shown in Figure 6 and discussed further below).

**Figure 5:**
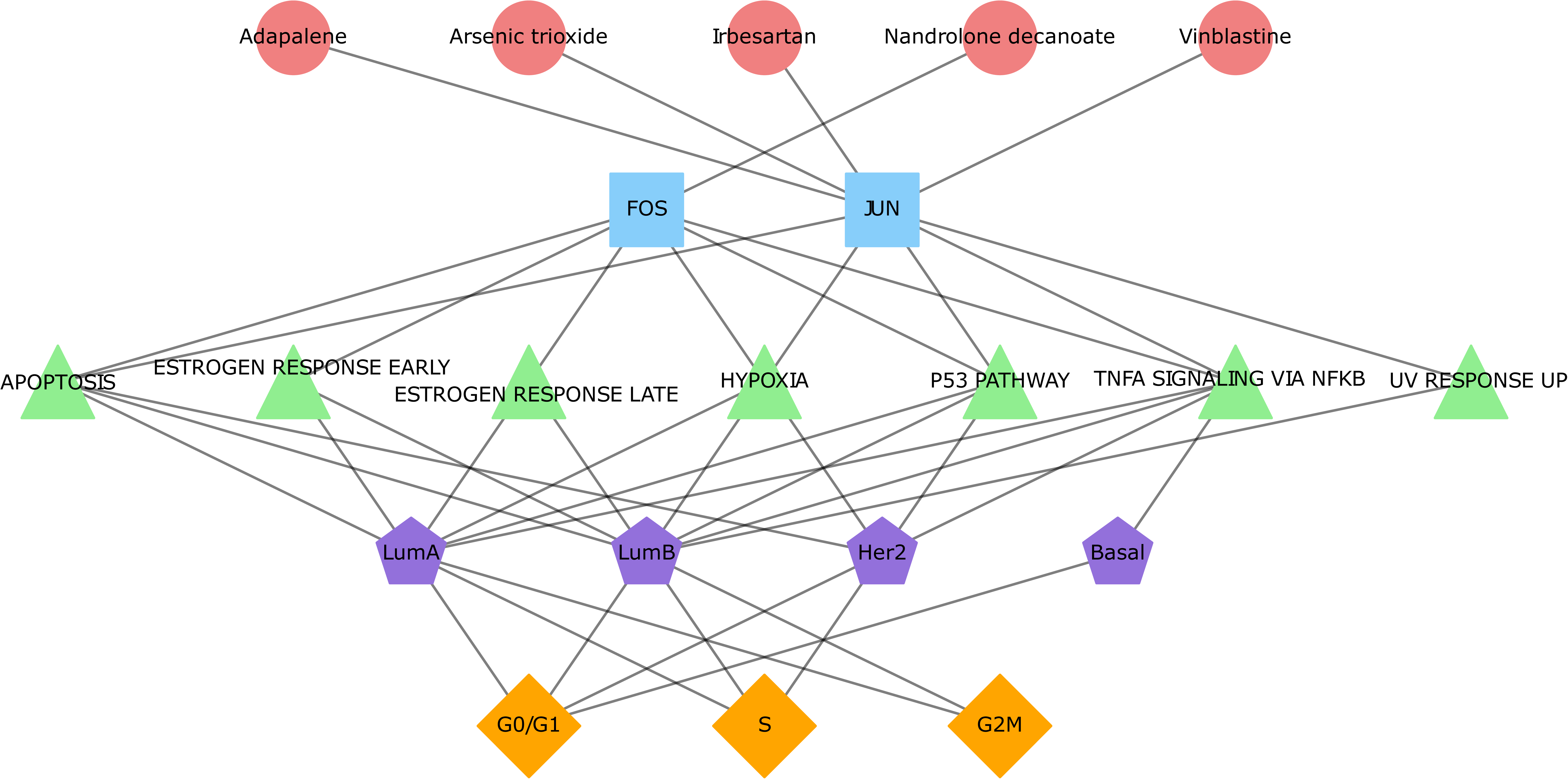
A multipartite network with five distinct levels: FDA approved drugs from Drugbank, transcription factor target genes, target hallmark pathways, breast cancer subtypes, and cell cycle phases. Associations are represented as links.

**Figure 6:**
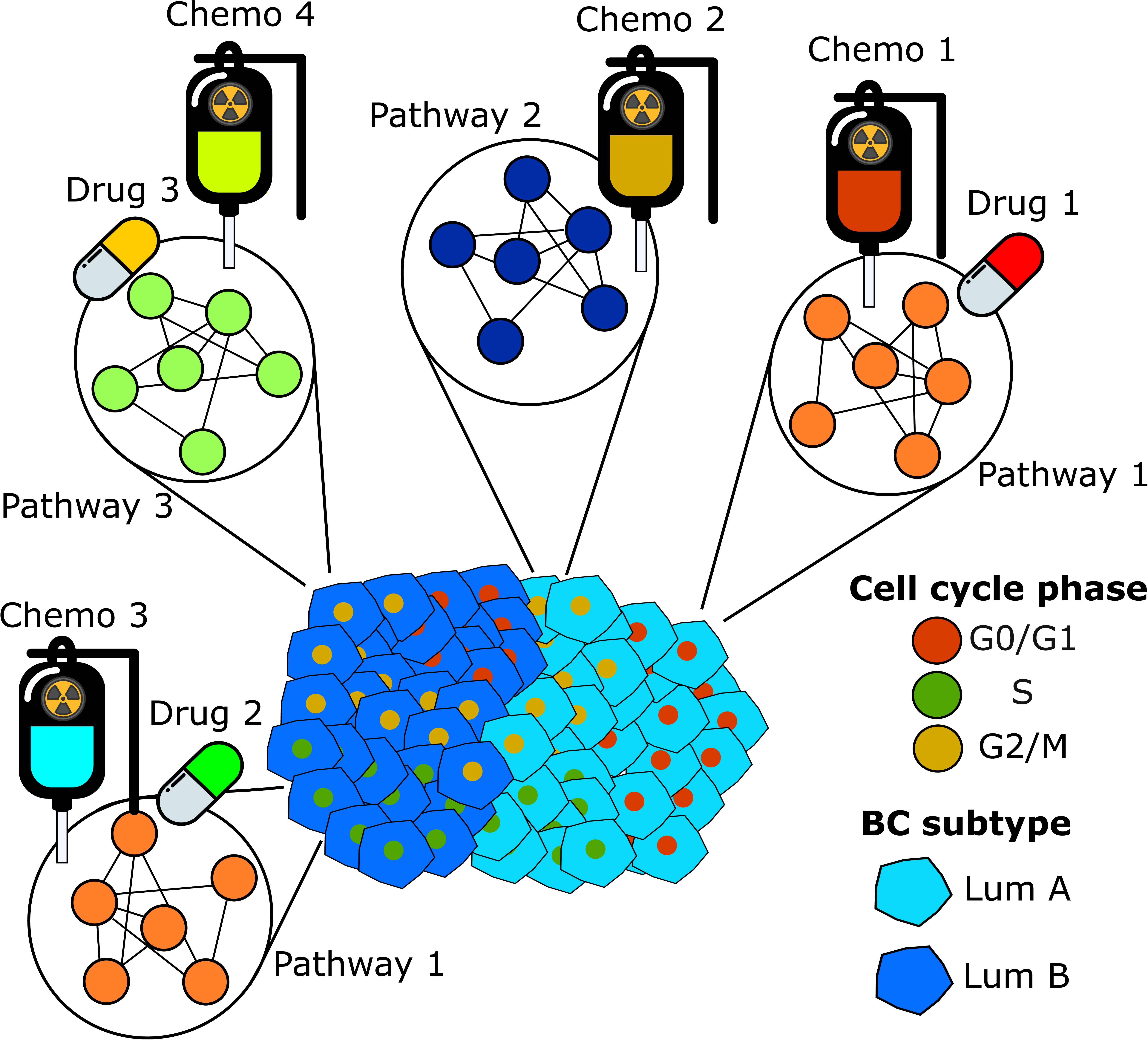
Example of potential applications of subtype- and cell cycle phase-specific therapies for a heterogeneous breast cancer tumor. Distinct tumor regions could be targeted with personalized combinations of drugs and chemotherapeutic agents, each selected based on the tumor’s breast cancer subtypes and specific cell cycle phases.

## Discussion

In this study, we examined the differences in cancer hallmark signaling pathways in cell cycle phases taking breast cancer subtypes into account. Our analysis involved two distinct approaches: comparing cells *across* different breast cancer subtypes that are in the same cell cycle phase and comparing cells *within* the same subtype that are in different cell cycle phases. In general, we found that taking cell cycle phases into account identifies a larger pool of relevant hallmark pathways that are not found when cell cycle phases are ignored. We showed that pathways that are enriched/depleted in one subtype when comparing *across* subtypes can also be enriched/depleted in one cell cycle phase relative to others when comparing *within* a single subtype (e.g. E2F_TARGETS in Luminal A). We also identified pathways that were uniquely enriched in specific subtype and cell cycle phases (e.g. MTORC1_SIGNALING and KRAS_SIGNALING_UP in G0/G1 HER2 and Basal cells, respectively). Finally, we provide examples of chemicals and FDA approved drugs that can target transcription factors upstream of relevant pathways e.g. the drug Vinblastine targets the transcription factor JUN which regulates a regulon whose genes significantly overlap with those in APOPTOSIS, P53_PATHWAY, TNFA_SIGNALING_VIA_NFKB and HYPOXIA, in G0/G1 phase Her2 cells.

Genomic analyses of the signaling pathways and cellular processes active in tumors have traditionally been performed on bulk samples. This means that the expression of genes across a broad range of heterogeneous cells is averaged and the signal coming from pathways that are differentially expressed in subgroups of cells can be diminished or lost. Recently, a better understanding of this heterogeneity has been facilitated through single-cell[5,30,43–45] and spatial[5,46–49] transcriptomic analyses. Building on this work, our analyses found both homogeneity and heterogeneity in the cancer hallmarks expressed between breast cancer subtypes and between cell cycle phases. Placing these findings in the context of published literature is challenging owing to previous studies having only compared hallmark pathways between bulk tumor molecular subtypes, however some notable parallels can be drawn. In general, we found that LumA and LumB cells showed similarity in depleted pathways relative to Her2 and/or Basal cells. Both the MYC_TARGETS_V1 and E2F_TARGETS hallmarks followed this pattern of depletion in LumA/B cells, in line with previous work from Schulze *et al.*[50] and Oshi *et al.*,[51,52] respectively. Conversely, the ESTROGEN_RESPONSE_EARLY and ESTROGEN_RESPONSE_LATE hallmarks were enriched in LumA and B cells relative to other subtypes, as expected given the hormonal-driven nature of this subtype[53]. Her2 cells showed enrichment of the OXIDATIVE_PHOSPHORYLATION hallmark, a result supported by recent work examining expression of an oxidative phosphorylation gene signature across breast cancer subtypes[51,52]. Similarly, for Basal cells, we found enrichment of the EPITHELIAL_MESENCHYMAL_TRANSITION and ALLOGRAFT_REJECTION hallmarks, both of which have been strongly linked to the Basal-like bulk tumor subtype[54] and tumors classified as TNBC by IHC[55], respectively. Heterogeneity was also observed between cell cycle phases within each subtype, with pathway changes aligning well with known biology. For example, the G2M_CHECKPOINT hallmark was depleted in G0/G1 and S and enriched in G2/M cells in all four subtypes and the MITOTIC_SPINDLE hallmark was depleted in G0/G1 and S and enriched in G2/M again, in all four subtypes except in LumA G0/G1 phase (Fig. 3c). Similarly, the MYC_TARGETS_V1 was depleted in G0/G1 and enriched in S and G2/M phases for Her2 and Basal cells matching its known role in regulating S phase entry[56,57]. These observations highlight the importance of considering both subtype-specific and cell cycle-dependent differences in cancer hallmark expression, laying the foundation for more refined analyses of potential therapeutic targets.

To place our results in a more treatment focused context we sought to identify drugs that could be candidates for targeting specific TF-subtype-phase-pathway combinations. Here we highlight pertinent previous published data for a selection of drugs and chemicals but note that an exhaustive review of all drugs found in our analyses is beyond the scope of this work. Among these drugs we found that Adapalene, a third-generation topical retinoid used to treat acne vulgaris, emerged as a target drug for hallmark pathways including HYPOXIA in Her2 cells in G0/G1 phase. Adapalene and other retinoids are notable as they have previously demonstrated promising anti-cancer potential and efficacy as therapeutic agents targeting breast cancer stem cells[58]. Similarly, both Vinblastine, which disrupts the mitotic spindle apparatus, causing cellular arrest during mitosis and inducing apoptosis[59] and Irbesartan, an angiotensin receptor blocker used to treat hypertension that was shown to reduce metastasis in hepatocellular carcinoma[60]. Hallmarks including APOPTOSIS and P53_PATHWAY were found to be potentially targetable in LumA and B cells in S phase using the androgen and anabolic steroid Nandrolone decanoate. This steroid has however been tested in a clinical trial setting in the 1980s and did not improve overall response rate in advanced breast cancer when added to Tamoxifen[61,62]. Whilst we only focused on FDA approved drugs in our results, we also found significant hits for chemicals in CTD. Acetylcysteine for example, was found to target ESTROGEN_RESPONSE_EARLY and ESTROGEN_RESPONSE_LATE in LumA and LumB cells in G0/G1 phase, and is used as a mucolytic in patients with lung conditions and to treat acetaminophen overdose and it has also demonstrated potential benefits in breast cancer[63–66]. These findings underscore the importance of considering not only subtype information but also cell cycle phases in drug targeting strategies. By integrating subtype-cell cycle-specific information, we can develop more effective, precision cancer medicine treatment approaches for breast cancer.

To showcase this potential application of our work, we propose a hypothetical situation where scRNA-seq analysis has been applied to a surgically removed breast tumor sample. We approach this scenario from an ideal future precision medicine perspective where treatments are only given on the basis of targets identified in tumor cells rather than the current standard of care[67,68]. As in our analysis above, scRNA-seq is used to first provide a detailed map of the tumor’s heterogeneity, showing that it contains cells of multiple molecular subtypes in different cell cycle phases (see illustration in Figure 6). Second, network analyses are used to derive specific subtype-phase enriched pathways where drug targets are identified. For this example, we focus on a tumor where the predominant single-cell subtypes are LumA and B (which is the most common case we saw in our two analyzed cohorts). A first option could utilize combination therapy, for review see[69]. Here, we could combine “Drug 1” and “Chemo 1” to target “Pathway 1” in LumA G0/G1 phase cells, “Chemo 2” to target “Pathway 2” in LumA G2/M phase cells, “Chemo 3” and “Drug 2” to target “Pathway 1” in LumB S phase cells, and “Drug 3” and “Chemo 4” to target “Pathway 3” in LumB G2/M cells (Figure 6). A second strategy could be to employ sequential therapy, where a single chemotherapeutic agent or drug is administered per treatment course in a sequential fashion. This method may be particularly beneficial when taking cell cycle dynamics into account by arresting cells in specific phases before administering the subsequent chemotherapeutic agent to potentially enhance therapeutic efficacy. Going back to the same illustrative example, we could start with “Chemo 2” to target “Pathway 2” in LumA G2/M phase cells; this would arrest cells in a specific phase, introducing a non-arbitrary distribution of cells into the remaining phases. For instance, if the “Chemo 2” arrests cells in G2/M phase, that would mean that cells will be in either G0/G1 or S phase, reducing the variability of cell cycle phases and thus, optimizing the usage of the next drug. Subsequently, for the next sequential drug, in the illustrative example, only the “Chemo 3” and “Drug 2” would be needed for LumB cells in S phase. In general combination therapy would be a more “bulk” treatment process where all relevant pathways could be targeted simultaneously, and sequential would allow for a finer grain treatment where a more specific selection of drugs/chemo agents would be used. However, the toxicities associated with combination therapies would likely preclude their usefulness in our hypothetical approach.

There are also some limitations to this study. First, despite the plethora of tools available to infer the cell cycle phase of each cell[70], these methods typically lack generalizability and do not capture cell cycle dynamics properly[19]. Acknowledging these challenges, we relied on Seurat for cell cycle phase inference as it has been benchmarked and shown to be an effective and computationally efficient approach that is suitable for large cohorts. Second, we analyzed the G0 and G1 cell cycle phases together due to the challenge of accurately distinguishing between these transcriptionally similar states[70]. New studies have emerged that may be better able to separate these two phases[71] however this methodology has not been benchmarked or tested against other computational methods. Third, this analysis represents a snapshot in time. Cells were analyzed in terms of “belonging” to a specific cell cycle, but these cells will actively progress through further phases. Despite this, we believe our results to be robust and representative of processes present during each phase - this is owing to the aforementioned methodology of combining cells across a broad number of tumors and using a second cohort to confirm all findings.

In conclusion, our study shows the importance of cell cycle phases in understanding breast cancer biology, increasing the biological insights compared to bulk analysis. The distinct landscape of biological mechanisms active in each subtype-phase specific scenario suggests

that drug candidates targeting these pathways may offer a more targeted and effective approach to breast cancer treatment.

## Funding

This work was supported by the Iris, Stig och Gerry Castenbäcks Stiftelse for cancer research (N.P.T.); the King Gustaf V Jubilee Foundation (N.P.T.); the Stockholm Cancer Society (Cancerföreningen i Stockholm to L.S.L.); the Swedish Cancer Society (Cancerfonden, N.P.T. grant number: 200802, 232670 Pj; L.S.L. grant number: 222081, 220552SIA, A.M. grant number 21 0277 JCIA 01 H); the Swedish Research Council (Vetenskapsrådet, grant number 2020-02466 to L.S.L); the Swedish Research Council for Health, Working life and Welfare, (FORTE, grant number 2019-00477 to L.S.L.); ALF medicine (grant number FoUI-974882 to L.S.L.) and the Gösta Milton Donation Fund (Stiftelsen Gösta Miltons donationsfond, to L.S.L).

## Authors’ contributions

MCA and NPT contributed to the study concept and design. MCA contributed to the acquisition and analyses of data. All authors interpreted the data, drafted the manuscript, read, and approved the final version.

## Ethics approval and consent to participate

Publicly available data - Not applicable

## Data availability

The data used in this study are two publicly available breast cancer atlases, Wu et al.[5] and Pal et al.[30].

Code to reproduce the results of this study is publicly available at https://bitbucket.org/tobingroup/bc_decoding_cellcycle

## Competing Interests

N.P.T. has performed consultations to Pfizer. Alexios Matikas reports consultation/speaker for Veracyte, Roche and Seagen (no personal fees) and research funding paid to institutions by MSD, AstraZeneca, Novartis and Veracyte.

**Supplementary table 1:** Number of cells in each sc-Breast cancer subtype and cell cycle phase for A) Discovery cohort and B) Validation cohort.

**Supplementary table 2:** FDA approved drugs from Drugbank and chemical candidates from the Comparative Toxicogenomics Database (CTD) that target specific combinations of transcription factors, pathways, breast cancer subtypes, and cell cycle phases.

**Supplementary figure 1:** A multipartite network with five distinct levels: Chemicals from Comparative Toxicogenomics Database (CTD), transcription factor target genes, target pathways, breast cancer subtypes, and cell cycle phases. Associations are represented as links.

## Abbreviations

IHC: Immunohistochemistry
ER: Estrogen receptor
PR: Progesterone receptor
HER2: Human epidermal growth factor receptor 2
scRNA-seq: Single cell RNA-sequencing
DGE: Differential gene expression
FGSEA: Fast Gene Set Enrichment Analysis
TF: Transcription factor
GRN: Gene regulatory network
RSS: Regulon Specificity Score
CTD: Comparative Toxicogenomics Database
FDA: Food and Drug Administration

## Notes

https://bitbucket.org/tobingroup/bc_decoding_cellcycle/src

