## Supplementary figures and images for "Decoding Cell Cycle Phase Variations in Cancer Hallmarks Across Breast Cancer Subgroups"

### Supplementary Figure 1

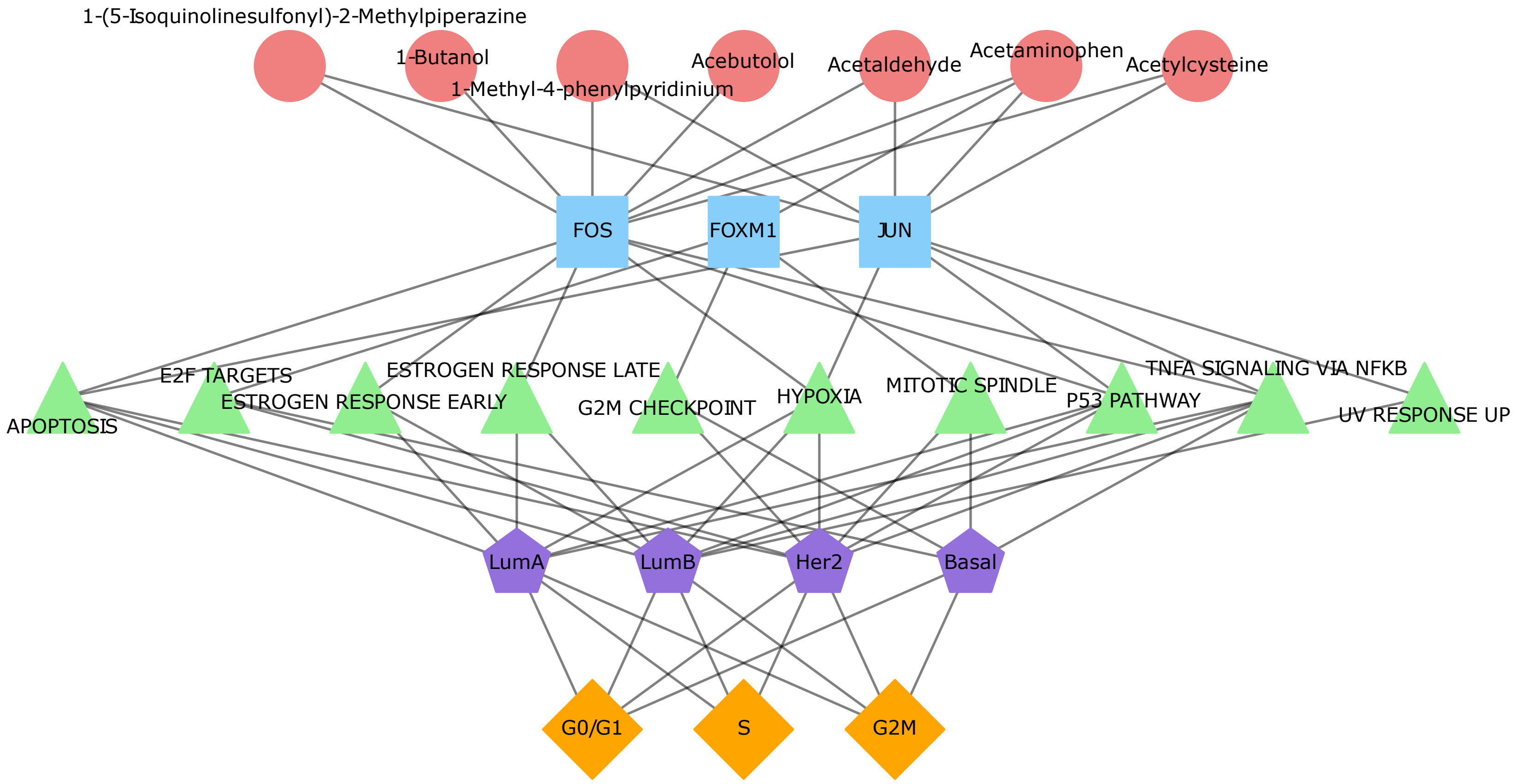
