## Supplementary Table 1 for "Decoding Cell Cycle Phase Variations in Cancer Hallmarks Across Breast Cancer Subgroups"

Discovery dataset

| **Breast cancer sc-subtype** | **Cell cycle phase** | | | |
| --- | --- | --- | --- | --- |
| **Calls** | **G0/G1** | **S** | **G2M** | **Total** |
| **Luminal A** | 3619 (53.2%) | 2435 (35.8%) | 755 (11.1%) | 6809 (27.8%) |
| **Luminal B** | 3214 (58.6%) | 1671 (30.5%) | 600 (10.9%) | 5485 (22.4%) |
| **Her2-enriched** | 3368 (51.7%) | 1942 (29.8%) | 1199 (18.4%) | 6509 (26.6%) |
| **Basal-like** | 2791 (49.1%) | 1612 (28.4%) | 1283 (22.6%) | 5686 (23.2%) |
| **Total** | 12992 (53.1%) | 7660 (31.3%) | 3837 (15.7%) | 24489 (100%) |
| Chi-square test p-value: <2e-16 | | | | |
| Cramer's V: 0.1 | | | | |

Validation dataset

| **Breast cancer sc-subtype** | **Cell cycle phase** | | | |
| --- | --- | --- | --- | --- |
| **Calls** | **G0/G1** | **S** | **G2M** | **Total** |
| **Luminal A** | 7122 (53.8%) | 3761 (28.4%) | 2364 (17.8%) | 13247 (13.9%) |
| **Luminal B** | 19598 (53.9%) | 10362 (28.5%) | 6418 (17.6%) | 36378 (38.1%) |
| **Her2-enriched** | 15529 (55.1%) | 7260 (25.8%) | 5369 (19.1%) | 28158 (29.5%) |
| **Basal-like** | 7887 (44.8%) | 5264 (29.9%) | 4467 (25.4%) | 17618 (18.5%) |
| **Total** | 50136 (52.6%) | 26647 (27.9%) | 18618 (19.5%) | 95401 (100%) |
| Chi-square test p-value: <2e-16 | | | | |
| Cramer's V: 0.062 | | | | |
